# The Lateral Habenula to Ventral Tegmental Area Pathway is Required for Aversive Learning and Defensive Behaviors

**DOI:** 10.1101/2025.08.28.672934

**Authors:** Marina R. Ihidoype, Jose Cesar Hernandez Silva, Kelly-Ann Pellerin, Ekaterina Martian, Maryse Pinel, Christophe D. Proulx

## Abstract

The lateral habenula (LHb) provides aversive signals to the ventral tegmental area (VTA), but its contribution to learning and behavior remains poorly understood. Using a retrograde viral strategy, we targeted VTA-projecting LHb neurons and monitored calcium activity during active avoidance training. These neurons were activated by aversive stimuli and predictive cues as animals acquired avoidance responses and showed increased activity at movement onset during the tail suspension test (TST). Silencing LHb→VTA transmission impaired avoidance learning, prolonged escape latency, and reduced the persistence and vigor of active coping in the TST, without affecting baseline locomotion. Anatomical and *ex vivo* electrophysiology revealed that LHb terminals innervate both dopaminergic (TH⁺) and non-dopaminergic (TH⁻) VTA neurons, exhibiting session-specific synaptic adaptations during avoidance learning. Together, these findings identify the LHb→VTA pathway as a source of aversive predicting signals required for the acquisition of avoidance behavior and the persistence of active coping in aversive context.

**Highlights:**

- LHb→VTA neurons are required for aversive learning and adaptive avoidance
- These neurons respond to aversive stimuli, predictive cues, and movement onset
- Silencing the pathway impairs avoidance learning and reduces active coping in aversive contexts
- LHb→VTA inputs innervate DA and non-DA neurons and undergo learning-related plasticity

## Introduction

Learning to predict and avoid harmful outcomes is essential for survival. The lateral habenula (LHb) plays a central role in processing aversive information and shaping behavioral responses to threats^1^. LHb neurons are excited by unexpected punishments and by neutral cues that predict them, thereby contributing to the formation of aversive associative memories such as avoidance learning^2–6^. In addition to signaling predictive value, LHb activity is required for organizing escape and avoidance behaviors, as shown by the disruption of defensive responses following LHb inhibition or suppression of specific inputs^2, 7^.

The ventral tegmental area (VTA) is a major target of the LHb and contains dopaminergic, GABAergic, and glutamatergic neurons involved in both reward and aversive processing^8–12^. Subsets of VTA neurons are activated by aversive stimuli and contribute to defensive behaviors and associative learning^13, 14^. Dopaminergic neurons can promote escape responses^15, 16^, glutamatergic neurons are required for defensive behaviors^17, 18^, and both dopaminergic and GABAergic neurons have been implicated in forming cue-outcome associations^16, 19–21^.

The LHb provides direct excitatory inputs to dopaminergic, GABAergic and glutamatergic neurons in the VTA^22, 23^. Rabies tracing experiments indicate that ∼ 5% of all VTA inputs arise from the LHb, identifying it as one of the main sources of aversive signals to this region^24, 25^. Optogenetic stimulation of VTA-projecting LHb neurons produces place aversion, indicating that these neurons carry aversive signals^26^. However, it is unknown how the LHb→VTA pathway is recruited during learning, whether it specifically encodes predictive cues, and if its activity is required for the execution of defensive responses.

Here, we tested the hypothesis that the LHb→VTA pathway encodes aversive predictive signals that are necessary for avoidance learning and the initiation of active coping responses. We combined viral tracing approaches, *in vivo* calcium imaging, pathway-specific silencing, anatomical mapping, and *ex vivo* electrophysiology. We found that VTA-projecting LHb neurons are recruited by aversive stimuli and predictive cues during avoidance learning, and that silencing this pathway impairs both the acquisition of avoidance behavior and active coping in an aversive context.

## Results

### VTA-projecting LHb neurons are activated by aversive stimuli and predictive cues during avoidance learning

To examine the role of VTA-projecting LHb neurons in aversive associative learning, we used fiber photometry to monitor their calcium activity in mice trained in an active avoidance task^27–29^. These neurons were targeted by injecting a retrograde AAV encoding Cre recombinase (retroAAV-Cre) into the VTA and a Cre-dependent AAV encoding the calcium sensor GCaMP7s (AAV-DIO-GCaMP7s) into the LHb. An optic fiber cannula was implanted over the LHb for calcium recording (Figure 1A, Figure S2A). To confirm labeling specificity, we performed a dual retrograde strategy with Cre/mCherry from the VTA and FlpO/eGFP from the RMTg, a structure located just caudal to the VTA that receives dense LHb inputs (Figure S1)^30, 31^. Across four mice (881 neurons), mCherry+ (VTA-projecting) and eGFP+ (RMTg-projecting) LHb populations were largely non-overlapping, consistent with previous reports^32, 33^. Only ∼7% of labeled neurons expressed both markers, confirming that our approach reliably and selectively targets VTA-projecting LHb neurons.

**Figure 1.**
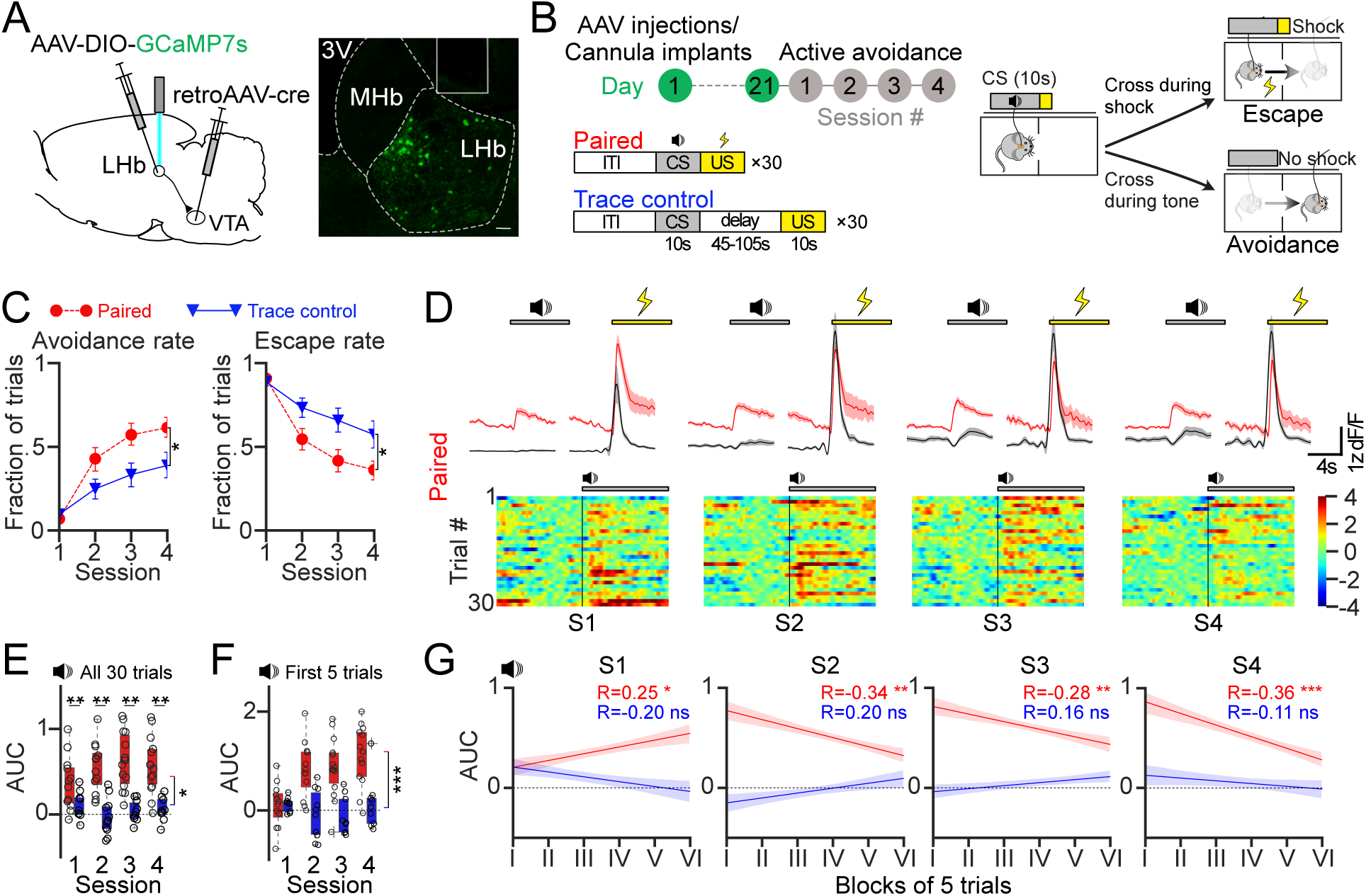
VTA-projecting LHb neurons are activated by aversive stimuli and predictive cues during avoidance learning. (A) Intersectional viral strategy: retroAAV-Cre was injected into the VTA and a Cre-dependent GCaMP7s AAV into the LHb, with representative confocal image showing GCaMP7s expression. (B) Experimental timeline and schematic of the active avoidance task under Paired (tone–shock association) and Trace control (tone with delayed shock) protocols. (C) Avoidance and escape rates across four sessions for Paired (red, n = 14) and Trace control (blue, n = 12) groups. (D) Peri-event plots of averaged calcium traces (z-scored ΔF/F) and mobility scores aligned to CS and US onsets, with single-trial heatmaps of CS-evoked activity across sessions (Paired group). The lines are means ± SEM. (E) Area under the curve (AUC) of CS-evoked calcium signals (0–2 s) across all trials of each session for Paired (red, n = 14) and Trace (blue, n = 12) mice. (F) Same analysis as in (E), restricted to the first 5 CS trials of each session. (G) Within-session dynamics of CS-evoked calcium responses, plotted as AUC in 5-trial bins. Data: mean ± SEM. Statistical test: linear mixed model (LMM), adjusted R values reported. ns, not significant; *p < 0.05; **p < 0.01; ***p < 0.001. See also Figure S1 and S2.

Mice were trained to associate a neutral conditioned stimulus (CS; tone) with a mild foot shock (US; 0.3 mA) (Figure 1B). In the Paired group, each trial began with a 10-second tone immediately followed by a 10-second foot shock. Crossings to the other compartment during the tone were scored as avoidances; crossings during the shock were scored as escapes. In the Trace control group, the CS and US were separated by a random 45–105 sec interval to disrupt associative learning while preserving exposure to both stimuli^34, 35^. Each session consisted of 30 trials, and mice were trained for four sessions. As expected, Paired mice rapidly increased avoidance responses across sessions, whereas avoidance remained significantly lower in the Trace controls (Figure 1C).

Mild foot shocks during the active avoidance and aversive airpuffs consistently elicited escape behavior (increased mobility scores) and evoked time-locked increases in calcium activity in VTA-projecting LHb neurons (Figure 1D, Figure S2B), consistent with their role in encoding aversive signals^26^. An auditory cue (tone) paired with foot shocks also evoked cue-driven activity during avoidance learning (Figure 1D-E).

Cue-evoked signals emerged within session 1: tones elicited little or no activity in early trials, but responses increased progressively during the session (Figure 1F–G). By sessions 2-4, cue-driven activity was present from the first trials and declined across trials, consistent with encoding aversive salience as mice avoid foot shocks (Figure 1F-G). These responses were absent in the Trace controls, confirming their dependence on CS-US contingency (Figure 1D-G, Figure S2C). In contrast, US-evoked responses remained stable across sessions, indicating consistent responses to aversive stimulus over time (Figure S2D).

### VTA-projecting LHb neurons activity increases at movement onset in both neutral and aversive contexts

We next examined whether activity in the LHb→VTA pathway was associated with movement onset independently of aversive stimuli. Calcium signals were recorded in the same mice used in the active avoidance task during the open field test (OFT), which measures general locomotor activity, and during the tail suspension test (TST), which evaluates motivated responses; both tests include alternating periods of mobility and immobility^28, 36^. Since general locomotor activity in the OFT did not differ between Trace control and Paired groups, data from both groups were pooled for calcium analysis (Figure S2E). Representative traces of calcium activity and corresponding mobility scores from individual mice are shown for both tasks (Figure 2A,D). In both the OFT and TST, calcium activity in VTA-projecting LHb neurons increased consistently at movement onset and decreased at mobility offset (Figure 2B-C and 2E-F). Calcium transients at movement onset, quantified as the area under the curve (AUC), were significantly larger in the TST than in the OFT (Figure 2G), indicating stronger recruitment of this pathway during movement initiation in an aversive context. Pearson correlation analysis further revealed a significant positive relationship between calcium activity and mobility scores in both tests, with correlation coefficients significantly higher in the TST (Figure 2H). This convergence suggests that LHb→VTA signaling is more strongly coupled to movement when behavior occurs under aversive conditions.

**Figure 2.**
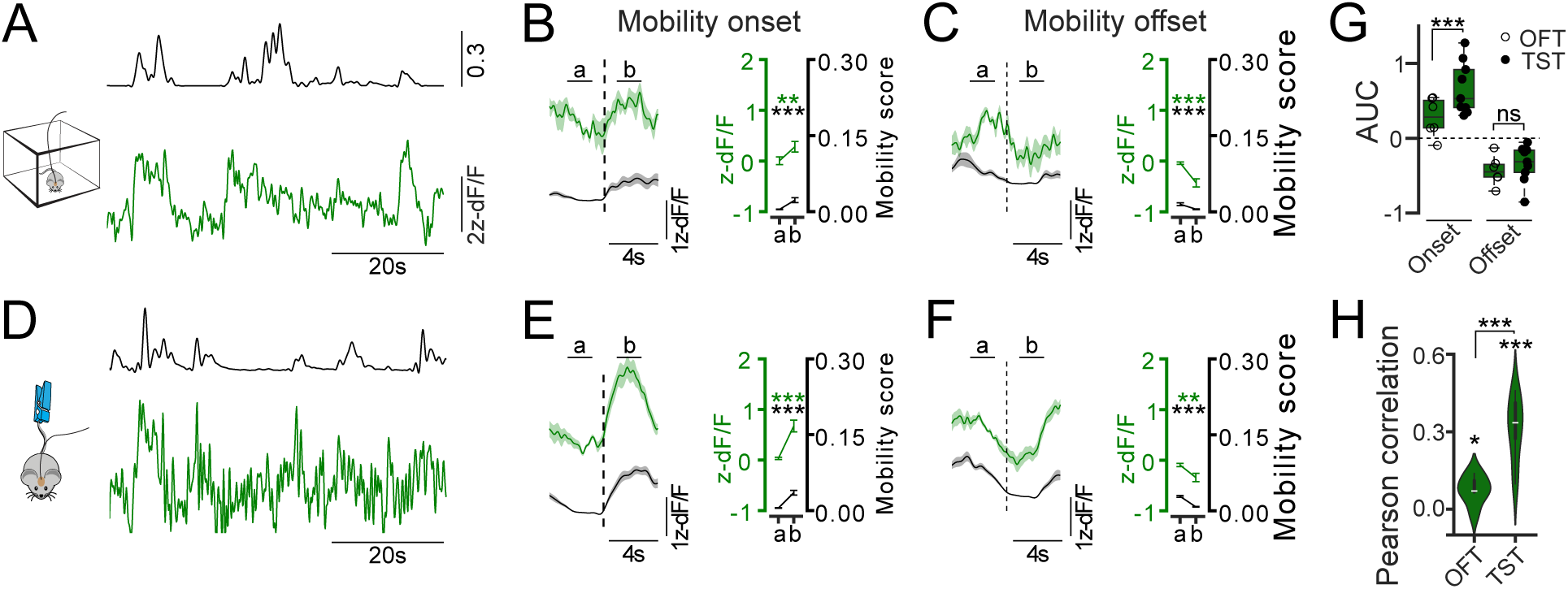
VTA-projecting LHb neurons activity increases at movement onset in both neutral and aversive contexts. (A) Representative calcium trace (z-scored ΔF/F) and corresponding mobility score during the open field test (OFT). B) Peri-event plots of average calcium activity and mobility score aligned to movement onset, with quantification of average responses in the 2 s window before (a) and after (b) onset (n = 6). (C) Peri-event plots of average calcium activity and mobility score aligned to movement offset, with quantification of average responses in the 2 s window before (a) and after (b) offset (n = 6). (D-F) Same convention as in (A-C), but during the tail suspension test (TST) (n = 9). (G) Area under the curve (AUC) of calcium activity in the 2 s window following movement onset and offset in the OFT (open circles, n = 6) and TST (filled circles, n = 9). (H) Pearson correlation between calcium activity and mobility scores in the OFT (n = 6) and TST (n = 9). Data: mean ± SEM. Statistical test: two-way ANOVA with post hoc Dunnett contrasts. ns, not significant; *p < 0.05; **p < 0.01; ***p < 0.001.

### Synaptic transmission in the LHb→VTA pathway is required for avoidance learning and the persistence of movement in aversive contexts

To determine whether the LHb→VTA pathway is necessary for associative learning and defensive responses, we constitutively silenced synaptic output from VTA-projecting LHb neurons. Using the same intersectional viral strategy, we expressed the tetanus toxin light chain (TeLC) fused to eYFP in VTA-projecting LHb neurons; control mice expressed eYFP alone (Figure 3A). TeLC cleaves the vesicle protein synaptobrevin, thereby blocking neurotransmitter release^37^.

**Figure 3.**
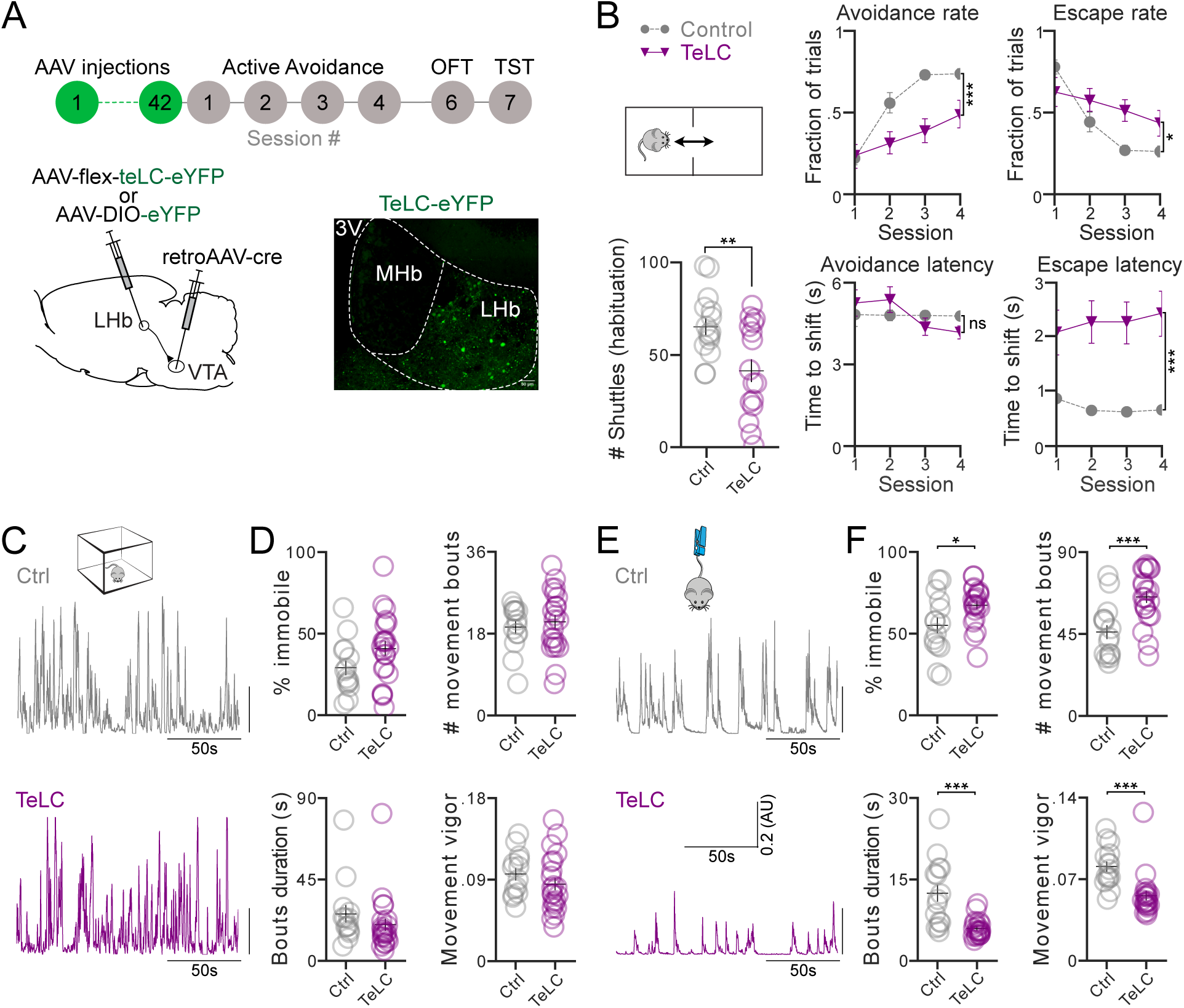
Synaptic transmission in the LHb→VTA pathway is required for avoidance learning and the persistence of movement in aversive contexts. (A) Experimental timeline and schematic of viral strategy for expression of TeLC-eYFP or control eYFP in VTA-projecting LHb neurons, with representative confocal image showing TeLC-eYFP expression in the LHb. (B) Group comparisons of baseline shuttles during habituation (session 1) and avoidance rate, avoidance latency, escape rate, and escape latency across four sessions of the active avoidance task for control (gray, n = 15) and TeLC (purple, n = 23) mice. (C) Representative mobility score traces for control and TeLC mice in the open field test (OFT). (D) Group comparisons of % time immobile, number of movement bouts, average bout duration, and movement vigor in the OFT. (E–F) Same convention as in (C–D) during the tail suspension test (TST). Statistical test: Welch’s t-test. ns, not significant; *p < 0.05; **p < 0.01; ***p < 0.001. See also Figure S3.

In the active avoidance task, TeLC mice made significantly fewer transitions between compartments during the 10-min habituation period of session 1 (Figure 3B). During training, they exhibited significantly fewer avoidance responses than controls across sessions 2–4 and showed longer escape latencies in all sessions (Figure 3B). Escape rates were also significantly reduced, whereas avoidance latency was unaffected.

Locomotor behavior was next assessed in the OFT and TST. In the OFT, TeLC mice did not differ from controls in immobility, number of movement bouts, bout duration, or movement vigor (Fig. 3C-D). Likewise, latency to move, total distance traveled, number of center entries, and speed in the periphery were unchanged (Figure S3A), confirming that general locomotor activity was intact. However, TeLC mice spent significantly more time in the center zone and moved more slowly within this zone (Figure S3A), consistent with reduced anxiety-like behavior.

In the TST, TeLC mice showed a distinct profile: they spent more time immobile, initiated a greater number of mobility bouts, but each bout was shorter in duration and lower in vigor (Figure 3E-F). Latency to move was unaffected (Figure S3B), indicating preserved initiation of movement. This suggests that the LHb→VTA pathway does not merely drive locomotion but encodes aversive salience that motivates defensive responses. Together, these results indicate that LHb→VTA transmission is dispensable for general locomotion but is required for movement persistence and vigor in aversive contexts, while its silencing also reduces anxiety-like behavior in the OFT.

### LHb terminals innervate dopaminergic and non-dopaminergic neurons in the VTA, with reciprocal connectivity

We used a dual-viral approach to identify VTA cell populations both innervated by LHb and projecting back to it. An AAV1-Cre virus, capable of anterograde transsynaptic transport, was injected into the LHb together with a Cre-dependent AAV encoding eGFP (AAV-DIO-eGFP) in the VTA to label VTA neurons receiving LHb input^38^. To identify VTA neurons projecting to the LHb, we injected a retrograde AAV encoding FlpO into the LHb and a FlpO-dependent mCherry reporter into the VTA. This strategy labeled LHb-innervated neurons (eGFP), LHb-projecting neurons (mCherry), and neurons with reciprocal connectivity (dual-labeled) (Figure S4A).

Quantitative analysis revealed that ∼50% of labeled VTA neurons were eGFP^+^, indicating they were innervated by LHb inputs, ∼25-29% were mCherry^+^, representing VTA neurons projecting to the LHb. And ∼20-25% were dual-labeled, indicating direct reciprocal connectivity between the LHb and VTA (Figure S4B-C). These findings align with prior studies describing reciprocal connectivity between the LHb and the VTA^24, 39, 40^. Immunostaining for tyrosine hydroxylase (TH) showed that 36-40% of LHb-innervated neurons were dopaminergic (TH^+^), and 60-64% were non-dopaminergic (TH^-^). TH^+^ targets were most common at intermediate VTA levels (bregma –3.16 to –3.28 mm), while TH^-^ targets were distributed across the rostrocaudal extent examined (–2.92 to –3.28 mm). These results confirm that LHb inputs to the VTA innervate both dopaminergic and non-dopaminergic populations, with a predominance of the latter, and that reciprocal LHb↔VTA connections are common. Given this mixed postsynaptic targeting, we next examined whether avoidance learning induces distinct synaptic adaptations at LHb terminals onto dopaminergic and non-dopaminergic VTA neurons.

### Avoidance learning induces population-specific plasticity at LHb terminals in the VTA

Given that LHb inputs innervate both dopaminergic (TH^+^) and non-dopaminergic (TH^-^) VTA neurons, we next examined whether avoidance learning drives population-specific synaptic changes at these synapses. We expressed AAV-ChR2-eYFP in the LHb and prepared acute VTA slices 1 h or 24 h after defined sessions of active avoidance learning under either Paired or Trace control protocols (Figure 4A). Optogenetic stimulation of LHb terminals was used to assess presynaptic release properties (paired-pulse ratio (P2/P1)) and postsynaptic strength (AMPAr/NMDAr ratio). Recorded neurons were filled with biocytin and immunostained post hoc for TH to distinguish TH^+^ and TH^-^ populations. In TH^-^ neurons, the AMPAr/NMDAr ratio was significantly increased in Trace control animals 24 h after a single training session, but not in paired animals or at other time points (Figure 4C). No significant AMPAr/NMDAr changes were detected in TH⁺ neurons under either protocol. In TH⁺ neurons, Paired training induced a significant decrease in P2/P1 ratio 1 h after the first training session, consistent with an increase in presynaptic release probability (Figure 4B). This effect was absent in the Trace control group and was not present at later training stages. In TH^-^ neurons, P2/P1 was almost significantly increased only after three sessions of Paired training, suggesting a shift toward reduced presynaptic release probability at later learning stages (Figure 4B).

**Figure 4.**
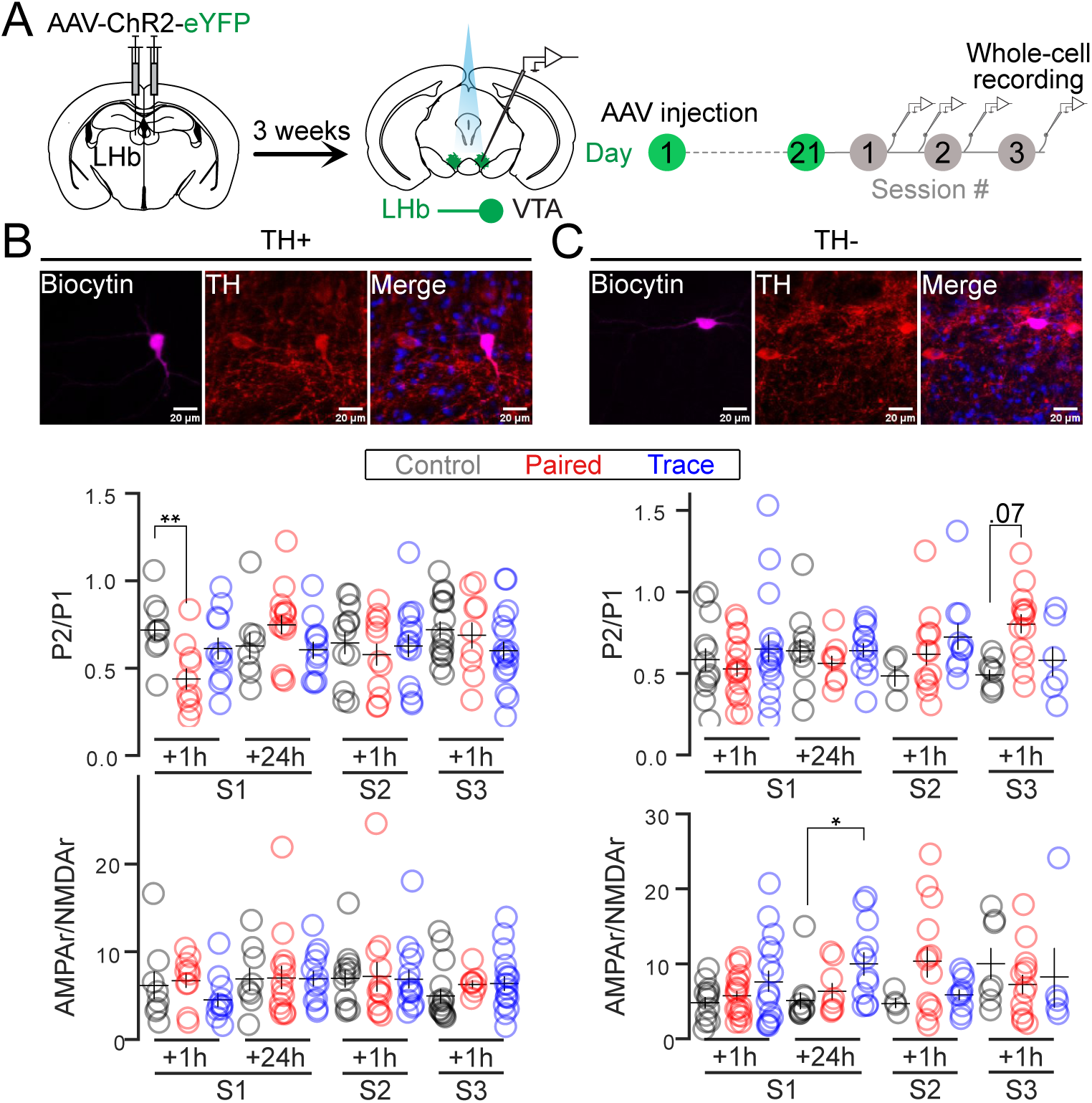
Avoidance learning induces population-specific plasticity at LHb terminals in the VTA. (A) Experimental design and timeline. AAV-ChR2-eYFP was injected into the LHb, and whole-cell patch-clamp recordings were performed in VTA neurons receiving LHb input. Recordings were obtained 1 h and 24 h after a single session of active avoidance, and 1 h after sessions 2 and 3, in Paired and Trace control protocols, as well as from naïve mice. (B) Confocal image of a VTA neuron filled with biocytin and positive for tyrosine hydroxylase (TH). Plots show paired-pulse ratios and AMPAr/NMDAr ratios measured 1 h and 24 h after session 1, and 1 h after sessions 2 and 3 in all groups. (C) Same convention as in (B), but for VTA neurons negative for TH. Statistical test: linear mixed model. *p < 0.05, **p < 0.01. See also Figure S4.

These results demonstrate that avoidance learning engages temporally distinct, cell-specific presynaptic adaptations within the LHb→VTA pathway, with TH⁺ neurons showing rapid early changes and TH⁻ neurons exhibiting delayed modifications after extended training.

## Discussion

The mammalian brain must detect threats and organize behavioral responses that maximize survival. Depending on the context, these responses may take the form of freezing to minimize detection or the initiation of active coping strategies such as escape or avoidance^15^. Our findings identify the LHb→VTA pathway as a critical circuit supporting both aversive associative learning and active coping behaviors.

Fiber photometry revealed that VTA-projecting LHb neurons are activated by aversive stimuli and by cues predicting them during avoidance learning. These neurons developed robust cue-evoked activity within the first training session, a signal absent in Trace controls, confirming its specificity for learned association. By sessions 2–4, cue responses appeared in early trials but declined within-session, consistent with the idea that once animals learn the US can be avoided, the aversive salience of the CS diminishes. Similar decreases in LHb cue responses have been reported during extinction when an aversive CS is no longer followed by the US^3^. Silencing this pathway impaired avoidance acquisition and increased escape latency, confirming its necessity for learning cue–outcome associations that drive adaptive behavior.

We also found that VTA-projecting LHb neurons are engaged at movement onset in both neutral (OFT) and aversive (TST) contexts, with larger activity changes at movement onset in the TST. Activity was correlated with mobility in both tests, but this relationship was significantly larger under an aversive condition. Consistently, silencing the pathway spared locomotion in the OFT, but selectively impaired active coping in the TST, reducing persistence and movement vigor. Together, these results suggest that the LHb→VTA pathway contributes to both initiation and persistence of movement, with its relative role depending on context. In active avoidance, reduced transitions and increased escape latency point to impaired initiation of goal-directed escape. In the TST, preserved bout initiation but reduced bout duration and vigor indicate a role in sustaining active coping. Photometry data support both interpretations, with increased activity at movement onset and positive correlations with mobility over time. Initiation and persistence may therefore represent complementary processes supported by aversion-related motivational signals within the LHb→VTA circuit. In addition, while silencing transmission in this pathway did not alter overall locomotion in the OFT, TeLC mice spent more time in the center zone where they moved more slowly, suggesting reduced anxiety-like behavior without impairing motor output. One possibility is that the LHb→VTA pathway conveys aversive salience signals, potentially via dopamine neuron activation, that help engage active defensive responses^16^.

Our anatomical and *ex vivo* electrophysiological data further demonstrate that LHb terminals innervate both dopaminergic and non-dopaminergic VTA neurons, with a bias toward the latter. Associative learning induced population-specific presynaptic adaptations: inputs to dopaminergic neurons showed rapid changes after a single session, whereas inputs to non-dopaminergic neurons adapted after extended training. These dynamics suggest that LHb→VTA inputs differentially regulate VTA subpopulations to support both the acquisition and refinement of aversive learning. This provides a potential mechanism by which LHb activity at cue and movement onset can shape downstream motivational and motor systems. It is tempting to speculate that in pathological conditions, unpredicted and unavoidable aversive outcomes may induce maladaptive synaptic adaptations in this pathway, contributing to cognitive and behavioral deficits observed in depressive disorders^41, 42^.

The LHb provides direct excitatory inputs to VTA^DA^, VTA^GABA^ and VTA^Glu^ neuronal populations^22, 23^. A subset of VTA^DA^ neurons is activated by aversive events^13, 14^, potentially encoding motivational salience and promoting escape behaviors^15, 16^. VTA^Glu^ neurons are also recruited by aversive stimuli, and their inhibition abolishes defensive responses^17, 18^. Together, these findings suggest complementary roles for VTA^DA^ and VTA^Glu^ neurons in encoding motivational salience. Our results further suggest that LHb inputs engage both populations to organize defensive behaviors. The VTA also contributes to associative learning, particularly when cues acquire predictive value for aversive outcomes^43^. Both VTA^DA^ and VTA^GABA^ neurons contribute to this process. Dopamine transmission to the prefrontal cortex (PFC) is critical for establishing new stimulus-outcome associations, potentially via LHb-driven recruitment of PFC-projecting VTA^DA^ neurons^21, 26, 44^. In parallel, phasic activation of VTA^GABA^ neurons projecting to the nucleus accumbens, driven by LHb inputs, may also contribute to associative learning^19, 20^.

These findings complement previous work showing that LHb projections to the GABAergic rostromedial tegmental nucleus (RMTg) suppress behavior by inhibiting dopamine neurons, favoring freezing or behavioral disengagement^15, 28, 45–48^. Increased activity of this pathway promotes passive coping.

In contrast, our results identify the LHb→VTA pathway as a parallel output that facilitates active coping. Its activity contributes to the acquisition of cue–outcome associations and supports both initiation and persistence of movement in aversive contexts, potentially modulated by inputs from the LHA^2, 5, 29^. This dual-function framework suggests that the LHb dynamically shifts between passive and active defensive strategies by recruiting distinct downstream targets depending on environmental demands ^4, 13, 49^. Such functional specialization of LHb outputs may underlie adaptive behavioral flexibility and, when dysregulated, contribute to maladaptive responses observed in affective disorders^1^.

### Limitations of the study

In this study, we used an intersectional viral approach to selectively express the calcium sensor GCaMP7s in VTA-projecting LHb neurons, allowing us to isolate the contribution of the LHb→VTA pathway in freely behaving mice. While powerful, fiber photometry measures bulk calcium dynamics and cannot resolve the activity of individual neurons or potential subpopulations of VTA-projecting LHb neurons. Our findings suggest the LHb◊VTA pathway supports learning and behavioral control, possibly via distinct LHb subpopulations targeting different VTA cell types^7^. We also constitutively silenced this pathway using TeLC. This approach was effective to test necessity, but may allow for compensatory adaptations that influence behavior, limiting interpretation about temporally precise contributions during learning and coping. In addition, while we confirmed innervation of both dopaminergic and non-dopaminergic VTA neurons, we did not directly test the contribution of these distinct VTA populations *in vivo*. Since these VTA populations likely project to different downstream brain regions, it is possible that learning-related plasticity occurs selectively within projection-specific subpopulations, which could not be resolved here. Finally, our experiments do not address how LHb inputs interact with other afferents to the VTA, such as those from the amygdala^50^, lateral hypothalamus^51^, or BNST^52^, which likely also contribute to shaping defensive responses. Future studies using cell-type–specific recordings and temporally precise manipulations will be essential to resolve these questions and to fully understand how LHb→VTA transmission supports aversive learning and active coping.

## Acknowledgments

We thank Maryse Pinel for her technical assistance. We thank Arturo Marroquín for his advice in statistical analysis. We also thank the CERVO Canadian Optogenetics and Vectorology Foundry Core Facility for producing the viral vectors. C.D.P. is supported by the Canadian Institutes of Health Research grant PJT169117 and the Natural Science and Engineering Research Council of Canada grant RGPIN-2025-04515, and receives Fonds de Recherche en Santé du Québec (FRQS) Junior-2 salary support. M.R.I. was supported by a doctoral training scholarship from FRQS and J.C.H.S. was supported by a Merit scholarship program for foreign students from the FRQS and a doctoral scholarship from CONACYT.

## Author contributions

M.R.I. and C.D.P. conceived, supervised and directed the study. M.R.I., J.C.H.S. and M.P performed the experiments. M.R.I., J.C.H.S. and K.A.P. performed the formal analysis of the data. M.R.I. and C.D.P. interpreted the data. M.R.I. and C.D.P. wrote and edited the manuscript.

## Inclusion and diversity

We support inclusive, diverse, and equitable conduct of research.

## STAR Methods

### Lead contact

Further information and requests for resources should be directed to the lead contact Christophe Proulx.

### Material availability

This study did not generate new unique reagents.

### Experimental model and subject details

Male and female C57BL/6 mice of 8-14 weeks used in this study were purchased from Charles River and allowed 1 week of acclimatation before experimentations. Mice were housed 4 per cage under standard conditions at their arrival (12 h light: 12 h dark cycle at 22-23°C and free access to food and water) and were used during the 12h light time. All experiments were conducted in accordance with the Canadian Guide for the Care and Use of Laboratory Animals guidelines and were approved by the Université Laval Animal Protection Committee.

### Stereotaxic surgeries

Surgeries were performed under aseptic conditions. Mice were injected with carprofen (10% of body weight), anesthetized with isoflurane (5%), and then placed in a small animal stereotaxic apparatus where the skull was exposed by scissor incision as described before^29^. Viral titers ranged from 10¹² to 10¹³ genome copies per milliliter. Either 150 nl (for single-virus injections per area) or 200 nl (for dual-virus injections per area) was injected into the LHb, VTA or RMTg.

#### Viral injections

*Fiber photometry*: AAV2/9-CAG-FLEX-jGCaMP7s-WPRE was injected into the LHb, and AAV2/retro-CAG-CRE-P2A-mNeptune was injected into the VTA.

*Tetanus toxin Light Chain (TeLC) Silencing*: AAV2/5-hSyn-flex-TeLC-P2A-eYFP or control AAV2/9-Ef1a-DIO-eYFP was bilaterally injected into the LHb, and AAV2/retro-CAG-CRE-P2A-mNeptune was bilaterally injected into the VTA.

*Anatomical Tracing*: A mixture of AAV2/1-hSyn-cre and AAV2/retro-CAG-Flpo-T2A-TagBFP2 was injected into the LHb, and a mixture of AAV2/5-hSYN-DIO-eGFP and AAV2/5-Ef1a-fDIO-mCherry was injected into the VTA.

*Electrophysiology.* AAV2/8-hSyn-hChR2(H134R)-eYFP was bilaterally injected into the LHb for optogenetic activation.

All viruses were purchased from either the CNP Viral Vector Core at the CERVO Research Center or Addgene.

Injections coordinates (from Bregma):

*LHb*: AP -1.65, ML ±0.45, DV -2.85

*VTA*: AP -3.3, ML ±0.5, DV -4.6.

*RMTg*: AP -4.0, ML ±0.5, DV -4.2.

Fiber photometry Cannula Placement:

For fiber photometry experiments, a 400μm optic fiber cannula was unilaterally implanted just above the LHb (AP -1.65, ML ±0.45, DV -2.5). The side of implantation was alternated between mice.

### Fiber photometry

Calcium signals were recorded using either a custom-build system^27, 29^ or commercially available systems from Doric Lenses (Quebec City, CA) or Neurophotometrics (San Diego, US). Excitation light from two LEDs (415 nm and 470 nm) was focused through a 20x objective (numerical aperture, NA 0.39) onto an optical patch cord connected to the implanted cannula in the freely moving mice. The 415 nm wavelength served as an isosbestic control to correct for motion artifacts, while 470 nm excitation was used to detect calcium-dependent fluorescence. Emitted fluorescence was collected through the same patch cord and recorded by the system. Light intensity was adjusted to 50-70 μW at the tip for each wavelength. Excitation alternated between the two wavelengths at a total sampling frequency of 40Hz (20Hz per LED).

### *Ex vivo* slices electrophysiology

#### Slice preparation

Mice were deeply anesthetized with isoflurane and intracardially perfused with ice-cold NMDG artificial cerebrospinal fluid (aCSF) containing the following (in mM): 1.25 NaH2PO4-H2O, 2.5 KCl, 10 MgCl2-6H2O, 20 HEPES, 0.5 CaCl2-2H2O, 28 NaHCO3, 8 glucose, 5 Na-ascorbate, 3 Na-pyruvate, 2 thiourea, and 93 NMDG [osmolarity adjusted with sucrose to 300–310 mOsmol/L and pH adjusted to 7.3 with HCl 10N (oxygenated with 95%O2/5%CO2)^53^. Kynurenic acid (2 mM) was added to the perfusion solution on the day of the experiment. Coronal slices (250 µm) were then cut with a vibratome (VT2000; Leica) to obtain sections containing VTA. Slices were placed in a 32°C oxygenated NMDG-aCSF solution for 10 min before incubation for 1 h at room temperature (RT) in HEPES-aCSF solution containing the following (in mM): 1.25 NaH2PO4-H2O, 2.5 KCl, 10 MgCl2-6H2O, 20 HEPES, 0.5 CaCl2-2H2O, 28 NaHCO3, 2.5 D-glucose, 5 Na-ascorbate, 1 Na-pyruvate, 2 thiourea, 92 NaCl, and 20 sucrose (osmolarity adjusted to 300–310 mOsmol/L at pH 7.4).

#### Whole-cell patch-clamp electrophysiology

All data were acquired by using an Axopatch 200B amplifier, digitized with a Digidata 1500A, and analyzed with pCLAMP 10.6 software (Molecular Devices). Slices were transferred into the recording chamber of an upright microscope (Zeiss) and perfused at a rate of 3–4 ml/min with artificial cerebrospinal solution (aCSF) containing the following (in mM): 120 NaCl, 5 HEPES,2.5 KCl, 1.2 NaH2PO4, 2 MgCl2, 2 CaCl2, 2.5 glucose, 25 NaHCO3, and 7.5 sucrose. The aCSF in the perfusion chamber was kept at 32°C. For the recording of excitatory postsynaptic currents (EPSCs), gabazine (10 μM) was added to block inhibitory currents mediated by GABAA receptors. Cells were visualized with a 60× objective on an upright fluorescent microscope with a video camera (Zeiss).

Borosilicate glass patch pipettes (3–6MΩ) were filled with internal solution containing the following (in mM): 115 cesium methanesulfonate, 20 cesium chloride, 10 HEPES, 2.5 MgCl2,4Na2-ATP,0.4 Na-GTP,10 Na-phosphocreatine, 0.6 EGTA, 5 QX314, and 0.2% biocytin (pH 7.35). Signals were filtered at 5 kHz. Pipettes and cell capacitances were fully compensated. Optically evoked postsynaptic responses were obtained delivering 5 ms pulses of473 nm light through the light path of the microscope using a Colibri 7 LED light source (Zeiss) under computer control. Neurons were voltage-clamped at −60 mV to record AMPAr EPSCs and at +40 mV to record dual component EPSCs containing AMPAr and NMDAr EPSCs. At +40 mV voltage, EPSC from AMPAr and NMDAr are kinetically different. Consequently, to obtain the AMPAr/NMDAr ratio, the peak of the AMPAr EPSC was divided by the magnitude of the NMDAr EPSC measured 50 ms after stimulation, at this point AMPAr-mediated current is negligible^54^. This approach allowed us to sample a much larger cell population from an individual mouse. Paired-pulse ratios (PPR) were recorded by giving two 5 ms blue light pulses at −60 mV with a 100 ms interval and calculated by dividing the amplitude of the second peak by the amplitude of the first peak (averaged responses from 12 sweeps).

### Histology and immunostaining

#### After behavior or tracing

Mice were deeply anesthetized using a mix of ketamine/xylazine (100 and 10 mg/kg, respectively, intraperitoneally), and transcardially perfused with saline followed by a 4% w/v paraformaldehyde (PFA) solution. The brains were removed and post-fixated overnight in the same PFA solution, rinsed with phosphate buffer solution (PB), and stored in PB at 4°C until used for immunostaining. Brains were sliced along the coronal plane at 100 μm for histology and at 50 μm for immunostaining.

#### After electrophysiology

Following electrophysiology recordings, brain slices were incubated in PFA solution for 1 h at room temperature (RT), rinsed in 0.1 M PB solution (pH 7.4), and stored in PB at 4°C until used for immunohistochemistry.

#### Immunostaining

Free-floating coronal sections were first blocked in PB containing 3% normal serum (NS), 0.4% Triton X-100, and 0.3% bovine serum albumin (BSA) for 1.5 h at RT. They were then incubated overnight at 4°C with sheep anti-TH (1:500, MilliporeSigma AB1542) for VTA slices. Slices were then rinsed 3×10 min in PB and were incubated for 4h at RT with secondary antibodies diluted in PB containing 3% BSA and 0.4% Triton X-100. The secondary anti-body was donkey anti-sheep Alexa Fluor 647. DAPI was added to stain the nuclei and Streptavidin Alexa Fluor 488 conjugate (1:1,000, Thermo Fisher Scientific, S32354) to specifically label the neurons that were patched and filled with biocytin.

After rinsing, sections were mounted on slides using Fluoromount Aqueous Mounting Medium (MilliporeSigma, F4680). Fluorescent images were taken using a Zeiss LSM700 confocal microscope.

### Behavioral testing

#### Active Avoidance (AA)

Behavioral tests were performed during the light phase. The active avoidance (AA) task was performed using the two-compartment shuttle box (Med Associates), adapted for use in mice. To facilitate the movement of the fiber optic patch cords during the task, the door between compartments was inverted. Mice not undergoing testing were placed in a separate room to avoid pre-exposure to the auditory cue. Behavioral training consisted of up to four daily sessions (one session per day), each including 30 trials. Mice were randomly assigned to either a Paired or Trace control protocol. Session 1 began with a 10-min habituation period, during which mice could freely explore the shuttle box. Sessions 2 through 4 began with a 1-min habituation period.

In the Paired condition, each trial began with a 10-second auditory tone (36 dB, fixed frequency; conditioned stimulus, CS), immediately followed by a 10-second escapable foot shock (0.3 mA; unconditioned stimulus, US). Mice could terminate the foot shock by crossing to the opposite compartment. If the mouse crossed during the CS (before the shock onset), the trial was counted as an avoidance. If the crossing occurred during the US, the trial was counted as an escape. Trials with no crossing were scored as failures. The inter-trial interval (ITI) varied randomly between 20 and 40 seconds to minimize temporal prediction. To maintain consistency, the CS tone always originated from the compartment in which the mouse was located at trial onset. In the Trace control condition, the CS and US were temporally dissociated by a random delay (45–105 seconds) to prevent associative learning between the tone and the foot shock^34^.

#### Open Field Test (OFT)

Mice were placed in a white open field arena (50x50 cm) and allowed to freely explore for 10 minutes. Behavior was recorded using a camera from the top, and the animal’s position was tracked in real-time using a contrast-based method implemented in Bonsai software. For experiments involving fiber photometry, positional timestamps were aligned with the onset of calcium signal recordings to allow peri-event analysis of movement and neuronal activity.

#### Airpuff Test (AP)

Mice were placed in the same open-field arena (50x50 cm) and allowed to explore freely. During a 6-min session, five air puffs were delivered at variable intervals (50-70 s apart). Air puffs were administered using a handheld air can duster positioned approximatively 30 cm above the mouse. Each air puff was manually triggered and timestamped via key press, allowing alignment with calcium signals during post hoc analysis (see Martianova et al,. 2023)^29^. Behavior was tracked with the same strategy as in the OFT.

#### Tail Suspension Test (TST)

Mice were suspended by the tail for 20 minutes in a 3-wall closed cage, facing the back wall, not allowing them to see the environment. Behavioral tracking was performed using the same camera and Bonsai-based setup as in the open-field test, from the open side of the box by the side. Movement bouts were analyzed in relation to calcium dynamics where applicable.

### Data analysis

Fiber photometry data was analyzed as previously detailed^27, 29^. Briefly, calcium-dependent (*signals*) and calcium-independent (*references*) traces were smoothed using a band-pass filter (moving average was used in^27^), flattened by removing the baseline using an airPLS algorithm (adaptive iteratively reweighted Penalized Least Squares^55^), and standardized to a mean value of 0 and a standard deviation of 1.

*References* were fitted to *signals* using a non-negative robust least squares regression (Lasso algorithm), and *zdF/F* was calculated by subtraction from standardized *signals* and fitted *references*.

*zdF/F* was aligned and averaged around the specific events such as airpuff, mobility onset-offset, CS-US, for each animal across all trials. Not all animals and trials were included at this step of the analysis, only the ones with high signal to noise ratio, which were defined by experimenter.

To measure the change from a baseline, area under the curve (AUC) was calculated. For airpuff, baseline AUC was calculated at the time frames -2 – -1 s, airpuff at 0 – 1.5 s. For mobility onset, immobile AUC was calculated at the time frames -3 – -1 s, and mobility at 1 – 3 s. For mobility offset, mobile AUC was calculated at the time frames -3 – -1 s, and immobility at 1 – 3 s. In avoidance test, baseline AUC was calculated at the time frames -2 – 0 s and CS at 0 – 2 s. AUC was normalized to the duration of the time frames (*AUCnorm* = *AUC*/(*t1 − t0*), where t0 and t1 are the beginning and the end of the time frames respectively).

Similarly, mobility score was aligned and averaged around the events, and AUC was calculated. For correlation analysis between calcium signal and mobility score, traces were interpolated to the same frequency.

Data distributions were tested using the Shapiro-Wilk normality test. Parametric or nonparametric tests were chosen depending on the number of observations, the distribution, and the model. For most of the data, a multi-factor mixed ANOVA with post hoc Dunnett’s or Tukey tests was used. The exact tests are indicated in the figure legends. Significance was set at p < 0.05 in most tests. Electrophysiological and behavior experiments were replicated at least three times.

### Sample size

The sample size for each group for behavioral, anatomical, and electrophysiological experiments was determined from previously published work and from pilot experiments performed in our laboratory.

### Declaration of generative AI and AI-assisted technologies in the writing process

During the preparation of this work the author(s) used ChatGPT in order to improve language and readability. After using this tool/service, the author(s) reviewed and edited the content as needed and take(s) full responsibility for the content of the publication.

