## Supplemental Figures for "The Lateral Habenula to Ventral Tegmental Area Pathway is Required for Aversive Learning and Defensive Behaviors"

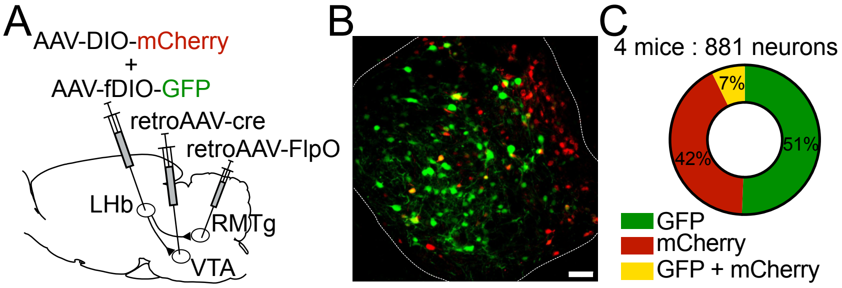


**Figure S1. Validation of selective targeting of VTA-projecting LHb neurons.**

(A) A retroAAV-Cre was injected into the VTA together with a Cre-dependent AAV encoding mCherry in the LHb, labeling VTA-projecting LHb neurons in red. In the same mice, a retroAAV-FlpO was injected into the RMTg together with a FlpO-dependent AAV encoding eGFP in the LHb, labeling RMTg-projecting LHb neurons in green.

(B) Representative confocal image of a coronal LHb section showing largely non-overlapping populations of VTA- and RMTg-projecting LHb neurons. Scale bar = 50 µm

(C) Quantification across four mice (881 neurons) revealed that 42% of labeled LHb neurons were VTA-projecting (mCherry+), 51% were RMTg-projecting (eGFP+), and only 7% were double-labeled.

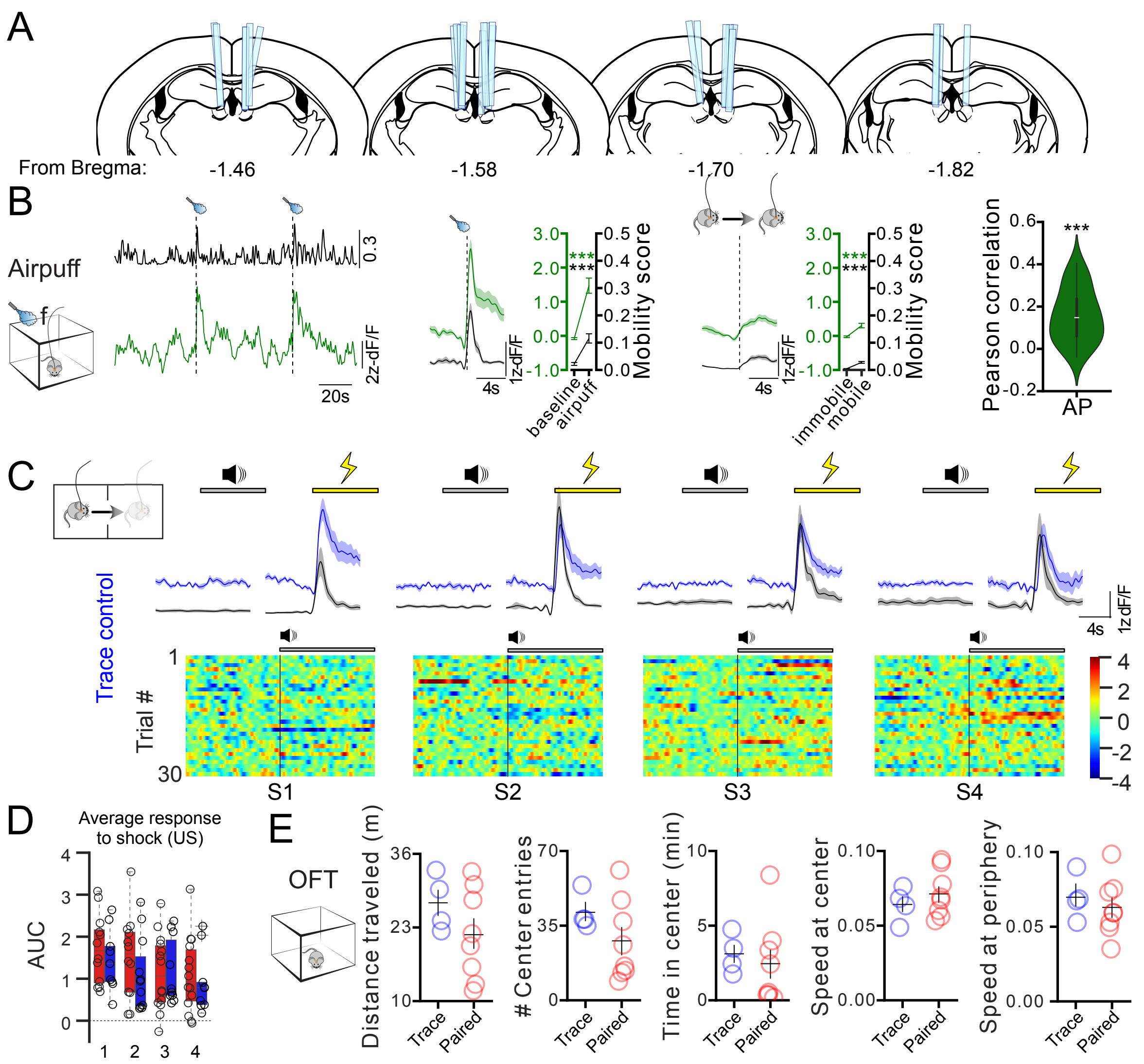


**Figure S2. Activity to aversive airpuff and active avoidance**

(A) Cannula placement for mice expressing GCaMP7s using in photometry recordings.

(B) Representative calcium trace (z-scored ΔF/F) and corresponding mobility score associated with the airpuffs (dashed vertical lines), Peri-event plot of the average calcium signal with all airpuff events at the VTA-projecting LHb neurons from Paired and Trace control mice (n = 12), plot of area under the curve before and after the airpuffs and Pearson correlation between calcium activity and mobility scores.

(C) Peri-event plots of averaged calcium traces (z-scored ΔF/F) and mobility scores aligned to CS and US onsets, with single-trial heatmaps of CS-evoked activity across sessions (Trace control group). The lines are means ± SEM.

(D) Area under the curve (AUC) of US-evoked calcium signals (0–2 s) across all trials of each session for Paired (red, n = 14) and Trace (blue, n = 12) mice.

(E) Group comparisons of distance traveled, # of center entries, time in center, speed at center and speed at periphery of Trace and Paired mice in the OFT.

Data: mean ± SEM. Statistical test: Linear mixed model (LMM), adjusted R values reported. ***p < 0.001. See also Figure 1 and Figure 2.

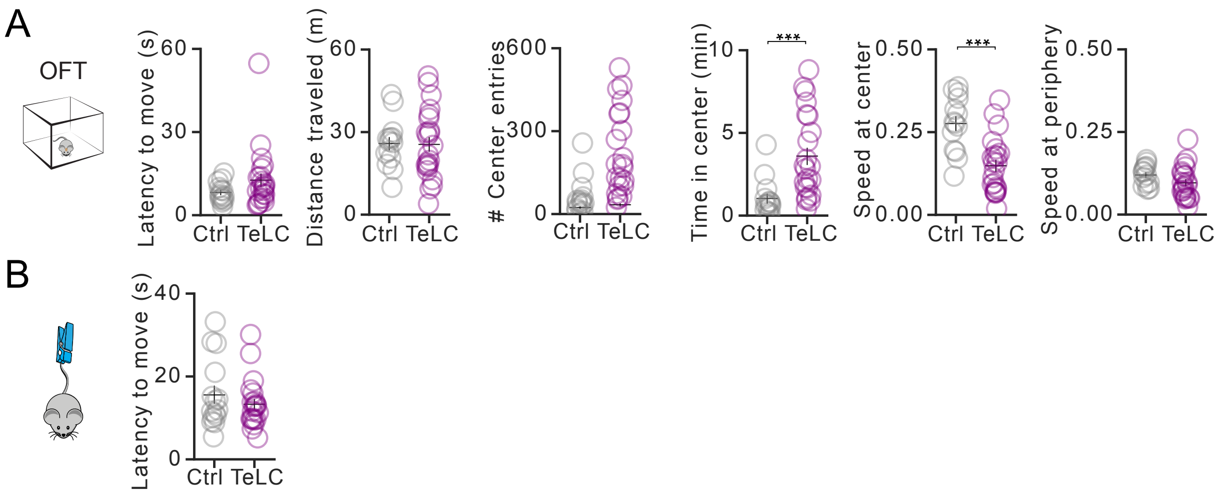


**Figure S3. Silencing transmission at the LHb→VTA pathway on behavior in OFT and TST.**
(A) Group comparisons of latency to move (s), distance traveled (m), # of center entries, time in center (min), speed at center and speed at periphery in the OFT.
(B) Latency to move (s) in the TST.
Statistical test: Welch's t-test. ***$p < 0.001$. See also Figure 3.

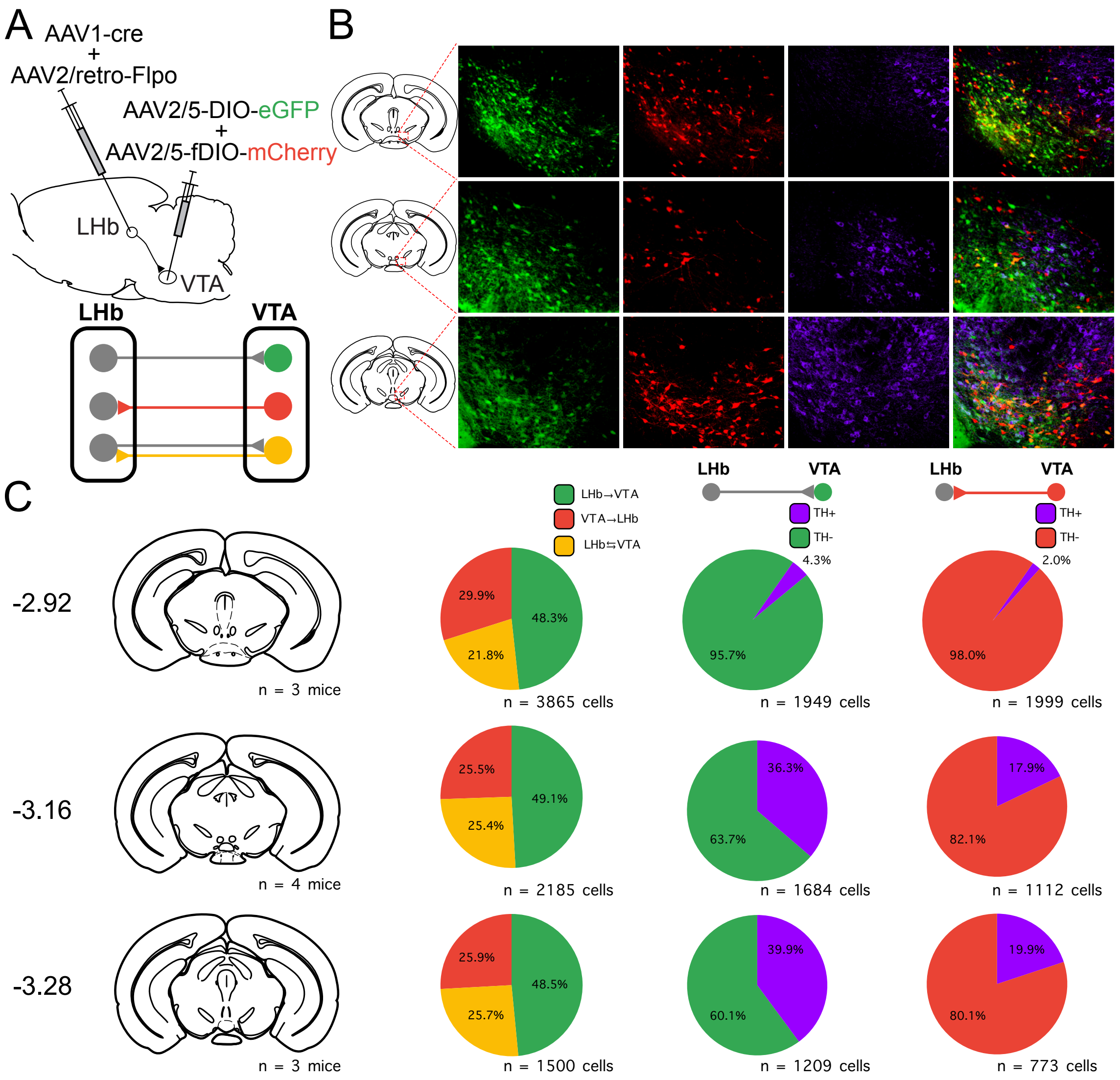


**Figure S4. Projections between the LHb and VTA are predominantly unidirectional and largely non-dopaminergic.**
(A) Viral tracing strategy using AAV1-Cre and AAV2/retro-FlpO injected in the LHb, combined with Cre- and Flp-dependent reporters (AAV2/5-DIO-eGFP and AAV2/5-fDIO-mCherry) injected in the VTA to label neurons projecting between LHb and VTA.
(B) Representative images showing eGFP, mCherry, and tyrosine hydroxylase (TH) expression in the VTA.
(C) Quantification of the fraction of labeled cells (eGFP$^+$, mCherry$^+$, or both) and their overlap with TH expression at multiple anteroposterior levels (relative to bregma). The majority of VTA→LHb and LHb→VTA projecting neurons do not express TH.
